# mTORC1-Dependent Regulation of the CCL24-CCR3 Axis Controls Granuloma Formation and Maintenance in Sarcoidosis

**DOI:** 10.1101/2025.07.11.664317

**Authors:** Xiongjian Rao, Jinpeng Liu, Derek B Allison, Douglas A Harrison, Ka Wing Fong, Yuanyuan Wu, Daheng He, Jia Peng, Zhiguo Li, Chi Wang, Jamie L. Sturgill, Parijat Sen, Xiaoqi Liu

## Abstract

Sarcoidosis is a chronic granulomatous disease marked by persistent inflammation and immune cell aggregation, yet its molecular underpinnings remain incompletely understood, hindering the development of effective targeted therapies. Here, we report that deletion of TSC1 or TSC2 in mice using a Fsp1-Cre leads to spontaneous formation of sarcoid-like granulomas, driven by hyperactivation of the mTORC1 pathway in fibroblasts and interstitial macrophages. Through inflammatory cytokine/chemokine array, we identified CCL24, a chemokine ligand for CCR3, as a key immunoregulatory molecule downregulated in both our murine model and sarcoid cohort plasma. Mechanistically, mTORC1 suppresses CCL24 expression via aberrant STAT3 signaling in fibroblasts and promotes CCR3 expression in interstitial macrophages, uncovering a novel regulatory axis in granuloma formation and maintenance. Pharmacological inhibition using rapamycin and azithromycin markedly attenuated granuloma burden and normalized CCL24-CCR3 signaling, underscoring the therapeutic relevance of this axis. Together, our study establishes a mechanistic link between mTORC1 activation, CCL24-CCR3 dysregulation, and granuloma persistence, offering not only a new insight into molecular mechanisms in sarcoidosis but also identifying promising targets for clinical intervention.

## Introduction

Sarcoidosis is a chronic inflammatory disease that can affect multiple organs in the body. Although it most commonly affects the lungs and lymph nodes, it can also affect other organs including the skin, eyes, liver, heart, and brain. The exact cause of sarcoidosis is unknown, but it is believed to result from overactive immune responses to unknown triggers, such as infections, genetic factors, chemical exposures, and/or stresses(Drent *et al*, 2021; Mukhopadhyay *et al*, 2012).

The hallmark of sarcoidosis is the formation of non-caseating granulomas, which are small clusters of immune cells that cause inflammation and can damage affected tissues(Costabel & Hunninghake, 1999; Grunewald *et al*, 2019; Krausgruber *et al*, 2023b). The pathogenesis of non-caseating granulomas may involve dysreglulated cellular composition and abnormal chemokine/cytokine production (Ng *et al*, 2000; Zhang *et al*, 2021). Though various biomarkers of sarcoidosis such as lysozyme (Tomita *et al*, 1999), angiotensin-converting enzyme (ACE) (Sahin *et al*, 2016) and more recently soluble IL-2R (Samiksha *et al*, 2022) has been studied, none have achieved clinical utility due to lack of a high positive predictive value and prognostic implications (Paolo *et al*, 2019). In addition, one of these molecules have provided potential targets for therapeutic discoveries in sarcoidosis.

The tuberous sclerosis complex (TSC)-mechanistic/mammalian target of rapamycin (mTOR) signaling pathway has been reported to regulate the innate inflammatory response (Weichhart *et al*, 2008; Zhu, 2014). TSC1 binds to and stabilizes TSC2, which is a GTPase activating protein (GAP) and downregulates mTORC1 activity through inhibiting small GTPase Rheb(Inoki *et al*, 2003; Saxton & Sabatini, 2017). It was reported that the deletion of TSC2 by Lyz2-Cre in mice causes granuloma formation, indicating that the activation of mTORC1 in macrophages is a pivotal hallmark of sarcoidosis (Linke, 2017). Additionally, it was shown that sarcoidosis granuloma formation is dependent upon mTORC1 transduction (Elliott *et al*, 2021).

In our study, studying the TSC1/2^flox/flox^/Fsp1 (fibroblast specific protein 1)-Cre mice, we found features consistent with sarcoidosis as well as identified the loss of CC chemokine ligand 24 (CCL24)/eotaxin-2, a key factor indicated in the pathogenesis of sarcoidosis (Meguro *et al*, 2020). Mechanistically, we found that mTORC1 suppresses CCL24 expression via aberrant STAT3 signaling in fibroblasts and promotes CCR3 expression in macrophages, uncovering a novel regulatory axis in granuloma formation and maintenance. Importantly, when we compared plasma of treatment naïve sarcoidosis patients with healthy human controls, we noted similar findings of lower CCL24 levels thus not only translating our KO mice as a reliable animal model for sarcoidosis but also raising the possibility of a new diagnostic biomarker in the form of CCL24.

Despite ongoing research, our current understanding of the underlying pathophysiology of sarcoid disease is limited. This is partially due to the lack of an effective diagnostic biomarker, which has also hampered the development of therapeutics. The current treatment modalities for sarcoidosis particularly corticosteroids, have severe side effects that affect the quality of life (Judson *et al*, 2015) and are not specific to sarcoid itself, but rather as general immunosuppressive agents (Grutters & Bosch, 2006). Thus, the development of more effective targeted therapies is a clinically unmet need.

Specific to the lung, azithromycin (AZM), a macrolide has had long found interest in lung disease both for its antibiotic and immunomodulatory properties(Fraser *et al*, 2020; Kournoutou & Dinos, 2022). Remarkably, in our mouse model, the administration of AZM and recombinant CCL24 effectively blocked the progression of granulomatous inflammation. This study reveals a critical role of CCL24 in the maintenance of sarcoidosis and improves critical and integrated understandings of the pathogenesis and development of sarcoidosis. Furthermore, this study provides a novel, putative biomarker of disease that may serve as a therapeutic target for the treatment of patients with sarcoidosis.

## Results

### TSC1/2 ^fl/fl^; Fsp1-cre mice show a sarcoid disease state including granuloma formation, inflammation and dysregulated hematopoiesis

When we selectively deleted the tuberous sclerosis complex (TSC) protein 1 using a fibroblast-specific cre recombinase (Fsp1-Cre) (Jackson Laboratory, strain # 027458) in mice, we found that the TSC1^flox/flox^ /Fsp1-cre^+^ mice (denoted as TSC1 Knockout (KO)) showed swollen feet and tails, as well as inflammation around the ear and neck, and occasionally hair loss within 6 months of age (Fig. EV1A). This phenotype started to emerge at the average age of 4-month-old, but some mice developed the phenotypes as early as 1.5-month-old. Since TSC1 protein stabilizes TSC2 and its deletion activates the mechanistic/mammalian target of rapamycin (mTOR) pathway (Henske *et al*, 2016), TSC2 floxed mice were crossed with Fsp1-cre^+^ mice to obtain TSC2 ^flox/flox^ /Fsp1-cre^+^ mice (denoted TSC2 KO) and assess phenotypic concordance. Results showed that the TSC2 KO mice also show the onset of swollen footpad and tails, abnormal femurs and tibias, inflammation around the neck and ear (Fig. EV1B), and occasionally ocular alopecia (Fig.EV1C), which is consistent with the development of alopecia reported in sarcoid patients (House *et al*, 2012). Furthermore, both TSC1 and TSC2 KO mice exhibited marked splenomegaly (Fig.EV1D) and reduced survival (Fig.EV1E).

To investigate the tissue edema in footpads and tails, we performed H&E staining and observed prominent non-necrotizing granulomatous inflammation, accompanied by increased mast cell infiltration in the surrounding soft tissue (Fig. 1A). Consistent with their splenomegaly, KO spleens were largely replaced by abundant non-necrotizing granulomas (Fig. 1B). Similarly, KO livers displayed non-necrotizing granulomatous inflammation surrounding portal tracts and replacing most of the hepatic parenchyma (Fig. 1C). These non-necrotizing granulomatous phenotypes were similar in nature to those reported in sarcoidosis. We next examined the lungs and interestingly found that KO lungs showed interstitial lymphangitic distribution of non-necrotizing granulomatous inflammation around veins and bronchioles mimicking the peribronchovascular-lymphatic distribution of sarcoidosis granulomas in human lungs (Fig. 1D). These findings were very similar to previous murine models of sarcoidosis using TSC2 KO (Linke *et al*, 2017).

**Figure 1.**
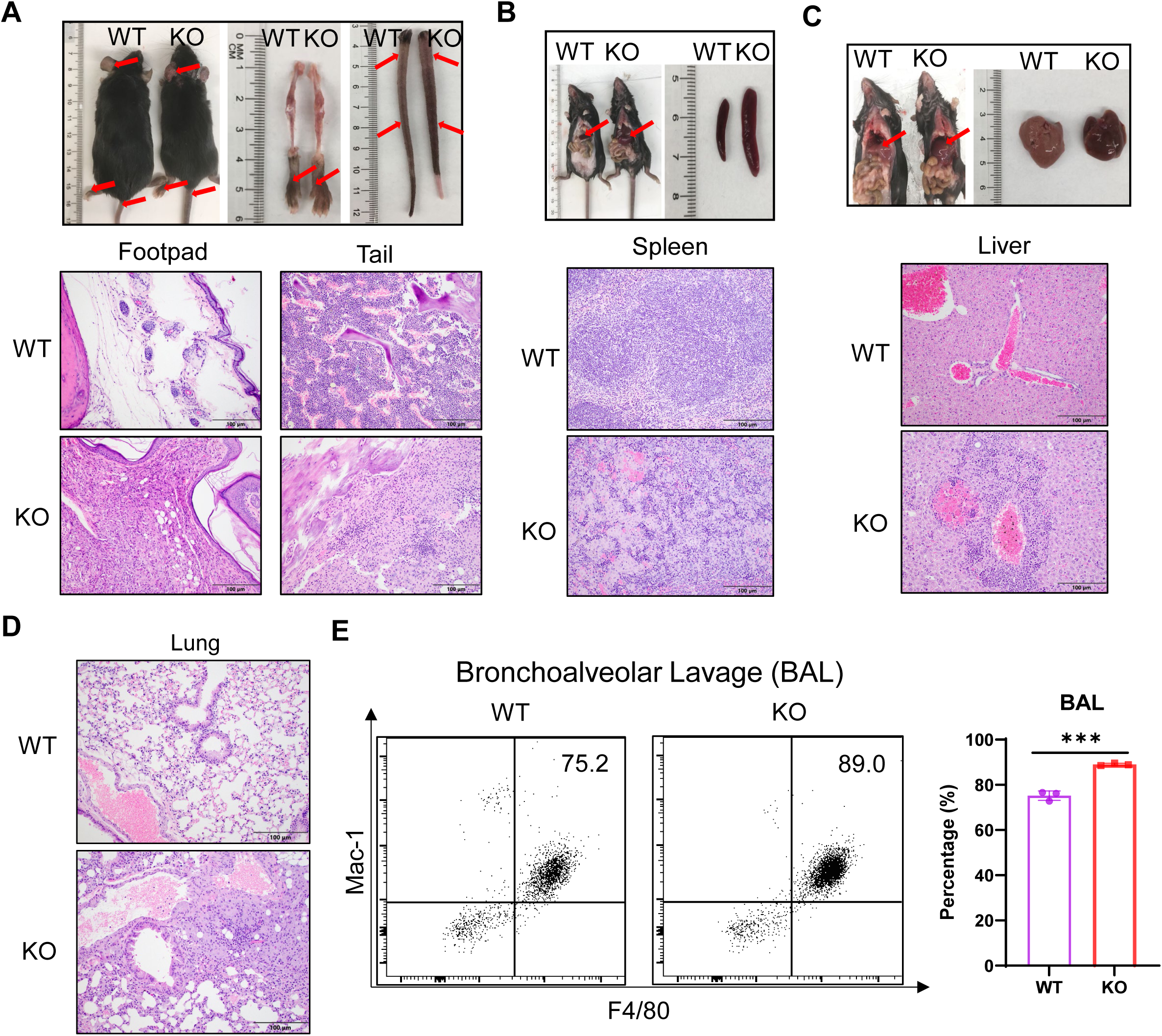
TSC1/2^fl/fl^; Fsp1-cre (TSC1/2 KO) mice show non-necrotizing granulomatous inflammation. H&E staining for the footpad as well as tail (**A**), spleen (**B**), liver (**C**), and lung (**D**) of the WT and KO mice. Scale bar,100µm. Results are representative of at least 6 mice per genotype. (**E**) The proportions of polarized macrophages in the bronchoalveolar lavage (BAL) from WT and KO mice. Data was analyzed by unpaired Student t test. Data are shown as means ± s.e.m. *** p<0.001.

Given sarcoidosis is predominantly driven by activated macrophages, we next examined macrophages from these KO mice by flow cytometry of bronchoalveolar lavage (BAL) samples. We found that Mac-1^+^F4/80^+^polarized macrophages were significantly increased in KO mice (Fig. 1E). Polarized macrophages were also markedly elevated in the peripheral blood, bone marrow, and spleen (Fig. EV2A). Given the increased mast cell infiltration in KO footpads and tails, we further analyzed peritoneal fluid and found a significant upregulation of mast cells (Fig. EV2B). Eosinophils (Joanne *et al*, 2014) were also significantly elevated in the lung (Fig. EV2C) and modestly increased in the liver (Fig. EV2D). These findings again corroborated with a sarcoid-like disease state of this mouse model.

Because the Cre utilized in this model is FSP1 and known to be expressed in fibroblasts, we investigated how the deficiency of TSC1 or TSC2 by this Cre affects hematopoiesis. Murine blood was collected analyzed by immunophenotyping flow cytometry. Results (Fig. 2A, and Table EV1) showed that KO mice had more white blood cells (WBC), especially significantly in TSC2 KO mice, and less red blood cells (RBC), indicating anemia, a feature also observed in human sarcoidosis (Lower *et al*, 1988). The WBC expansion was primarily driven by increased myeloid populations, including neutrophils, monocytes, and eosinophils, while lymphocyte counts were decreased (Fig. 2A). Analysis of lymphocyte subsets revealed a significant increase in CD4^+^CD8^-^ T helper cells in the KO bone marrow and thymus, but a decrease in the spleen (Fig. 2B), which again aligned with findings in human sarcoidosis (Shen *et al*, 2016). These cells have been implicated in granuloma formation (Co *et al*, 2004; Krausgruber *et al*., 2023b). B220^+^B cells were significantly reduced in the KO spleen and liver, but unchanged in the bone marrow (Fig. 2C). Given the more severe inflammatory phenotype in TSC2 KO mice, we further profiled their bone marrow and identified a significant expansion of CD115^+^Gr1^+^ inflammatory monocytes (Fig. 2D), consistent with chronic inflammation. Mac1^+^Gr1^+^ myeloid cells were also significantly elevated in the peripheral blood, bone marrow, and spleen of TSC2 KO mice (Fig. 2E). Histological examination of bone marrow confirmed disrupted hematopoietic architecture in KO mice (Fig. 2F). To dissect changes in both lineage and progenitor cells of hematopoiesis, we performed flow cytometry for bone marrow as well as spleen and found that lineage-negative (lin^-^)sca1^+^c-kit^+^(LSK) cells in the bone marrow of TSC2 KO mice were significantly decreased (Fig. 2G). The increased myeloid cells which were aforementioned in the KO mice are due to a higher percentage of granulocyte-monocyte progenitors (GMP) (Fig. 2H).

**Figure 2.**
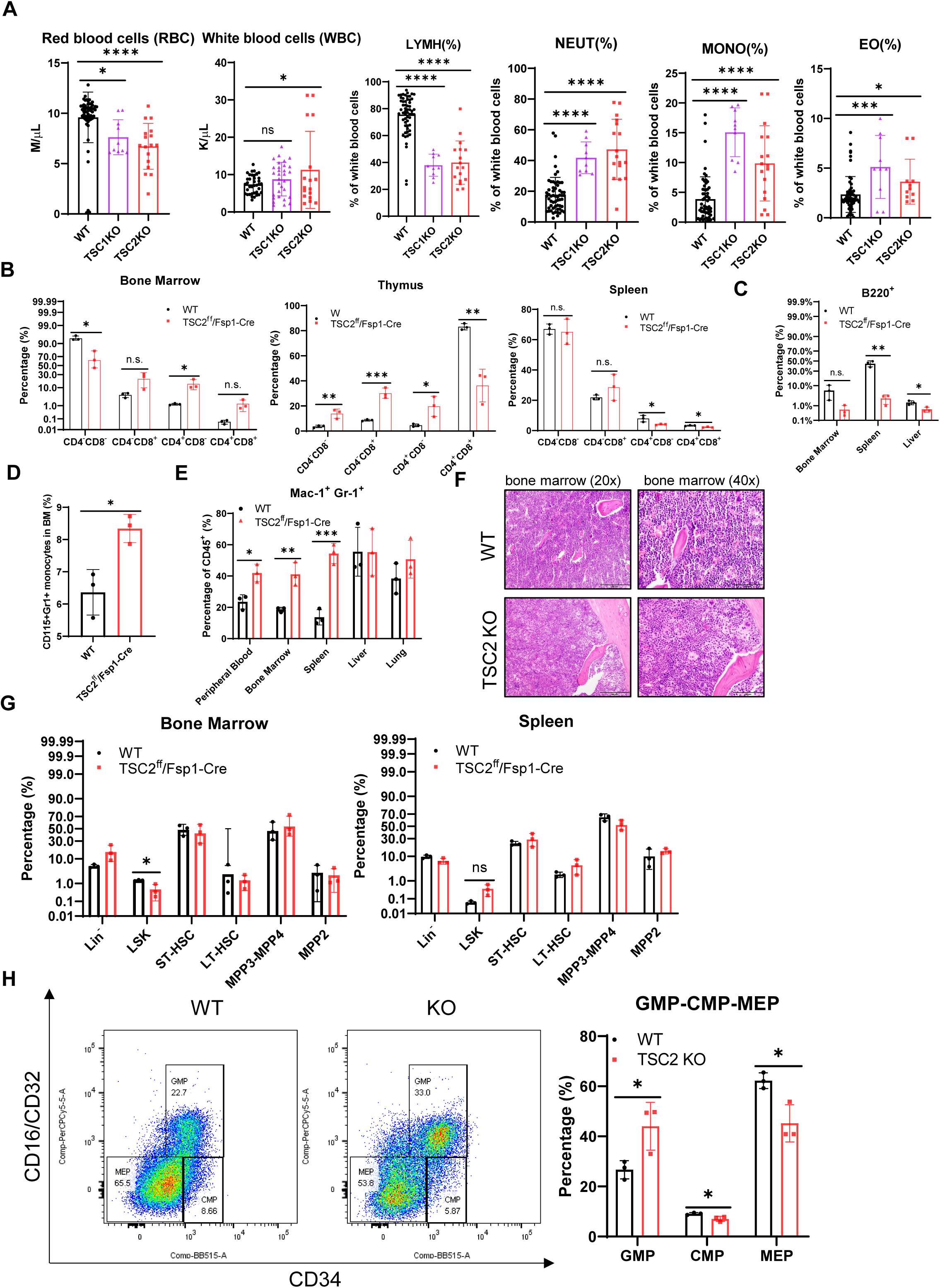
Hematopoiesis is dysregulated in TSC1/2 KO mice. **(A)** The dysregulated hematopoiesis by blood cell counting. **(B)** The proportions of CD4^+^, CD8^+^ T cells in bone marrow, thymus and spleen. **(C)** The percentages of B cells in bone marrow, spleen and liver. **(D)** The percentage of CD115^+^Gr1^+^ monocytes in bone marrow. **(E)** The percentages of Mac-1^+^Gr-1^+^ myeloid cells in peripheral blood, bone marrow, spleen, liver and lung. **(F)** H&E staining for the WT and TSC2 KO bone marrow. Scale bar,100µm. Results are representative of at least 3 mice per genotype. **(G)** The proportions of lineage cells, hematopoietic stem cells and progenitor cells in the bone marrow and spleen. The proportions of common myeloid progenitors (CMP), granulocyte-monocyte progenitors (GMP), and megakaryocyte-erythrocyte progenitors (MEP) in bone marrow. Data was analyzed by unpaired Student t-test. Data are shown as means ± s.e.m. * p<0.05, ** p<0.01, *** p<0.001, **** p<0.0001. n.s., not significant.

In brief, TSC1/2 ^fl/fl^; Fsp1-cre mice represent the sarcoidosis mouse model with additional pro-inflammatory comorbidities.

### mTORC1 activation in fibroblasts, interstitial macrophages, and proliferative macrophages contributes to the sarcoid state of TSC1/2 ^fl/fl^; Fsp1-cre mice

To determine which immune cell populations, contribute to the sarcoid state in the context of TSC1/2 deletion, we performed single cell RNA sequencing (scRNA-seq) on pulmonary immune cells, which is a major sarcoid organ. An unbiased clustering analysis was performed on lung samples from WT and KO mice (Fig. 3A), revealing major immune populations. Uniform Manifold Approximation and Projection (UMAP) clustering identified distinct immune cell populations, including alveolar macrophages (Avo_Macro) (Macro, Kcnip4), interstitial macrophages (Int_Macro) (Mafb, Maf), proliferating macrophages (Macro_prol) (Mki67, Top2a), CD8 T cells (Cd3e,Lck,Cd8a), CD4 T cells (Cd4), dendritic cells (DC) (Siglech,Cd300c, Kik1, Cd207), natural killer cells (NK) (Kirb1a, Nkg7, Gzma), regulatory T cells (Treg) (Ikzf2, Ctla4), and Neutrophils (Neu) (S100a9, S100a8, Retnlg, Camp) (Fig. 3B,C). Cell composition analysis revealed striking shifts between WT and KO lungs: alveolar macrophages (58.7%) dominated in WT lungs, whereas neutrophils (72.0%) and interstitial macrophages (12.3%) predominated in KO lungs (Fig. 3D). We then examined the gene expression level of TSC1 for Avo_Macro, Int_Macro, Macro_prol, CD8T, DC, NK, Treg, Neu, and found that TSC1 was markedly decreased in interstitial macrophages and proliferative macrophages from KO mouse lungs (Fig. 3E), which means that TSC1/2 deletion in fibroblasts has downstream effects in interstitial macrophages and proliferative macrophages of these mice and contributes to the pathogenesis of sarcoidosis.

**Figure 3.**
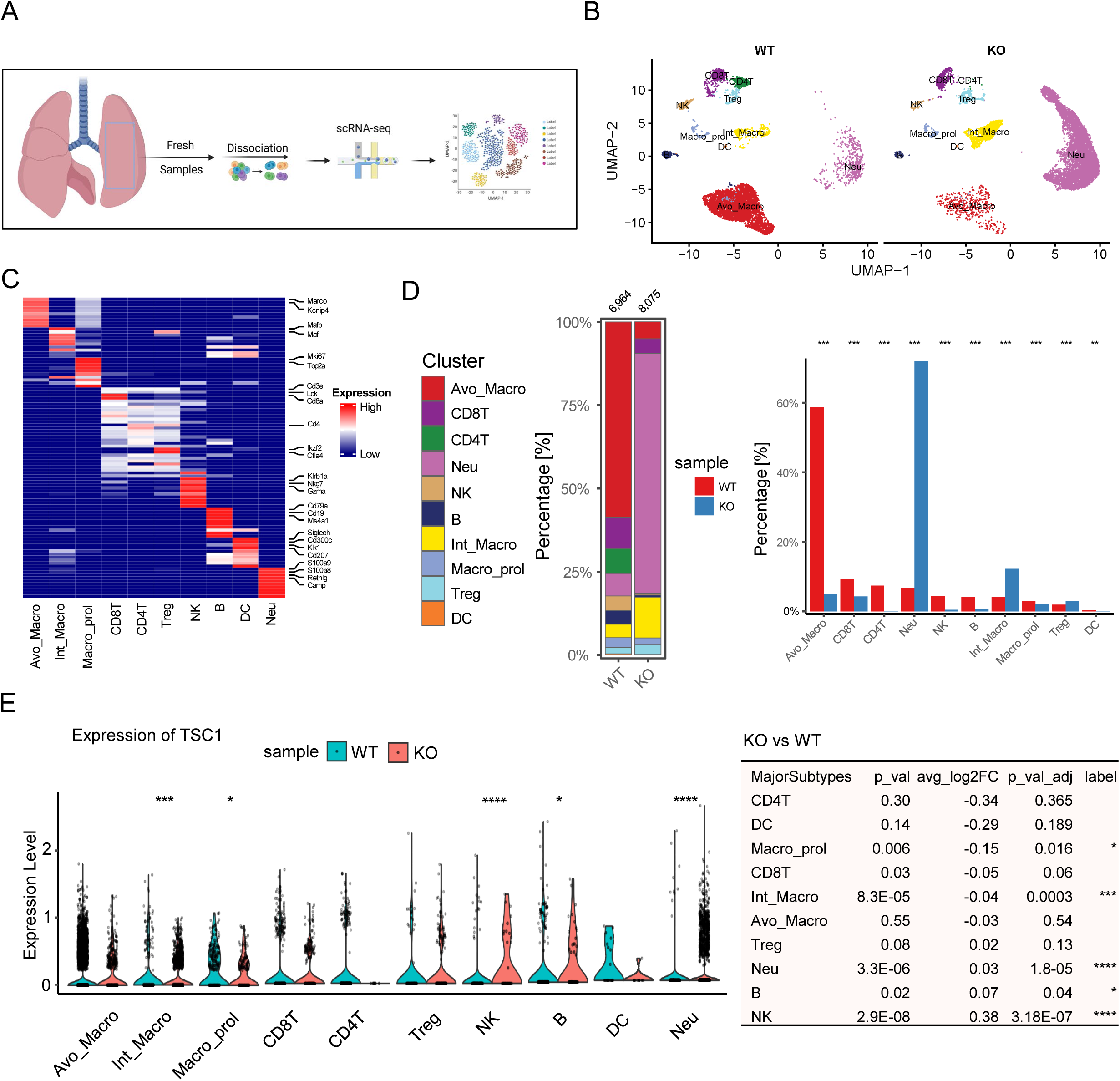
Single cell RNA sequencing for the lungs from the WT and KO mice. (**A**) The working model for the single cell RNA sequencing analysis. (**B**) The Uniform Manifold Approximation and Projection (UMAP) of cell types. (**C**) The heatmap of cell markers for each immune cells. (**D**) The proportions of different cell types among the total immune cells. (**E**) The expression level of TSC1 in major subtypes of immune cells. Data was analyzed by unpaired Student t test. Data are shown as means ± s.e.m. * p<0.05, ** p<0.01, *** p<0.001.

Single-cell analysis reveals transcriptomic remodeling in distinct cell types that contribute to the progression of sarcoidosis. To examine cell-cell communication underlying granuloma formation, we performed ligand-receptor interaction analysis using a curated database (Ramilowski et al., 2015). KO lungs exhibited a greater number and strength of inferred ligand–receptor interactions than WT lungs, indicating heightened intercellular communication (Fig. EV3A). In WT lungs, proliferating macrophages (Macro_prol) engaged most frequently in cellular crosstalk, whereas in KO lungs, interstitial macrophages (Int_Macro) and neutrophils (Neu) dominated the interaction networks (Fig. EV3B,C). Cytokines were categorized into four distinct clusters (Fig. EV3D) and signaling changes, both incoming and outgoing, were analyzed across key cell types including Neu (Fig. EV3E), Int_Macro (Fig. EV3F), CD4T (Fig. EV3G), Avo_Macro (Fig. EV3H). Notably, CCL chemokine signaling was among the most enriched signaling patterns in KO lung immune cells (Fig. EV3I), implicating CCL signaling in progression of pulmonary sarcoidosis. We further investigated the CCL signaling in different immune cells and found that CCL signaling was enhanced in Avo_Macro, Int_Macro, CD8T, CD4T and Treg cells from KO lungs (Fig. EV3J). Further analysis of CCL signaling revealed that cytokine distribution varies across immune cell types according to their abundance and function, underscoring the central role of cytokine networks in sustaining and advancing pulmonary sarcoid granulomas.

Collectively, these findings demonstrate that mTORC1 activation in fibroblasts, interstitial macrophages, and proliferative macrophages, together with dysregulated cytokine signaling, especially CCL signaling, drives the development and maintenance of sarcoidosis in TSC1/2 ^fl/fl^; Fsp1-cre mice.

### CCL24/eotaxin-2 is markedly downregulated in sarcoidosis and represents a potential diagnostic biomarker

To identify cytokine changes contributing to sarcoidosis in our model, we performed an inflammatory cytokine array using enzyme-linked immunosorbent assay (ELISA). Among the 40 cytokines and chemokines analyzed, CCL24 (also known as eotaxin-2) was dramatically reduced in the plasma of KO mice compared to age- and sex-matched WT controls (Fig. 4A, Fig. EV4A). In the bone marrow supernatant, CCL24 was again the most significantly downregulated cytokine, alongside changes in CCL11, TIMP-1, PF4, and leptin (Fig. EV4B). CCL24 downregulation in serum from KO mice was further validated by mouse CCL24 ELISA (Fig. 4B). Of significance, we found that CCL24 levels were significantly lower in the sera of human sarcoid patients compared to healthy individuals (Fig. 4C), consistent with findings in KO mice and supporting CCL24 as a potential diagnostic biomarker for pulmonary sarcoidosis. It is important to note these patients were all treatment naïve, thus excluding the possibility of treatment effects on lowered CCL24 expression. Given that TSC1/2 deletion in fibroblasts and fibroblasts are abundant, we assessed CCL24 expression in WT and KO murine embryonic fibroblasts (MEFs) and observed a marked reduction in KO MEFs (Fig. 4D). Treatment with the mTORC1 inhibitor rapamycin restored CCL24 expression in KO MEFs (Fig. 4E), linking its suppression to mTORC1 hyperactivation. To assess additional sources of CCL24, we evaluated its expression across hematopoietic cell types. CCL24 levels were significantly reduced in KO bone marrow, spleen, and bone marrow-derived mast cells (BMMCs), but not significantly changed in peritoneal fluid cells (PFCs) and bone marrow-derived macrophages (BMDMs) (Fig. 4F). Together, these results establish CCL24 downregulation as a consistent feature of sarcoidosis in both murine and human systems and support its potential utility as a diagnostic biomarker and mTORC1-regulated immune mediator.

**Figure 4.**
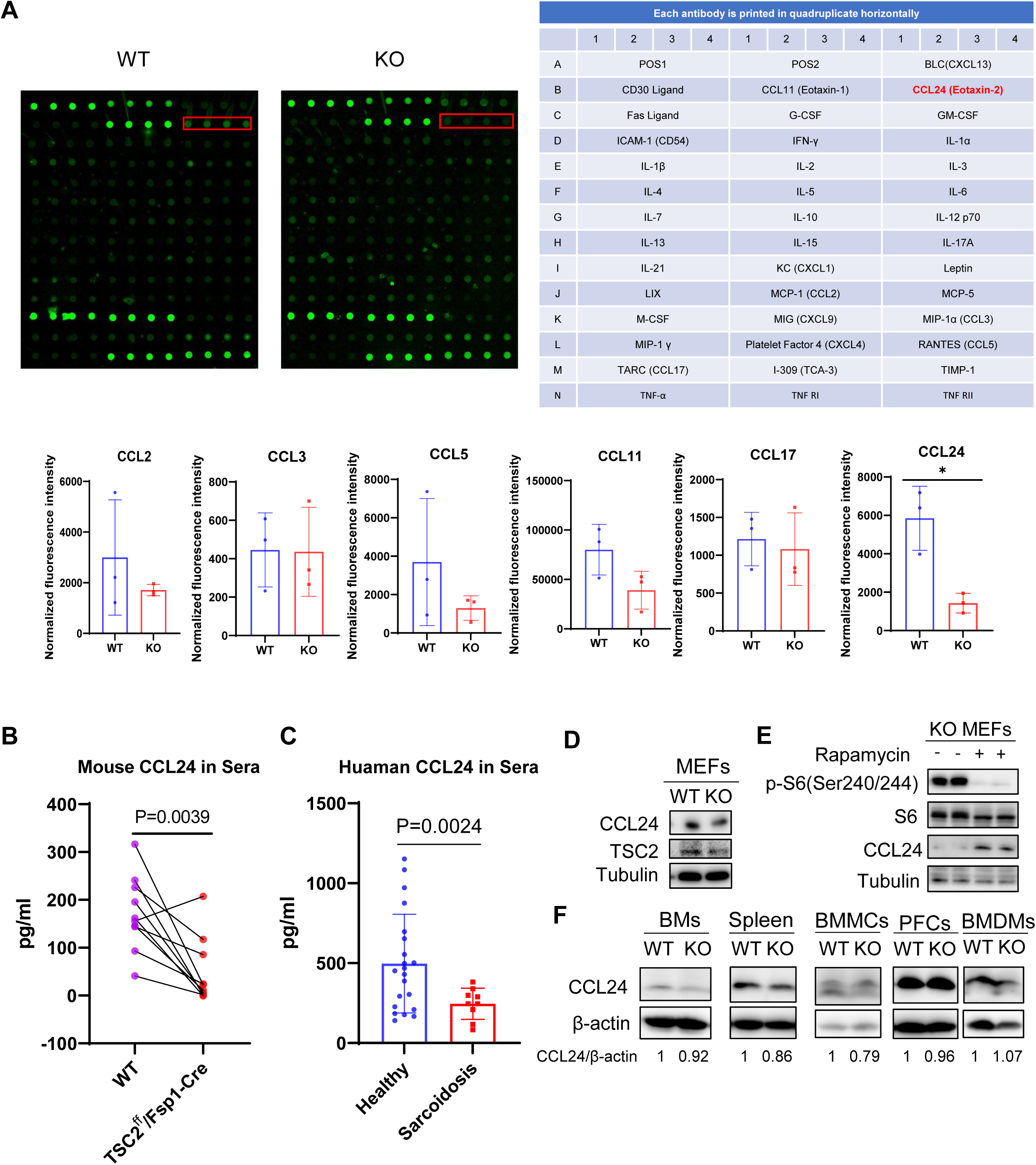
**CCL24/eotaxin-2 is decreased in TSC1/2 ^fl/fl^; Fsp1-cre mice. (**A) Inflammatory cytokine/chemokine array assay for blood plasma from WT and KO mice. (**B**) ELISA analysis for mouse CCL24 in blood plasms of WT and TSC2 KO mice. (**C**) ELISA analysis for human CCL24 in blood plasma of healthy human and sarcoid cohort. (**D**) Immunoblots of WT and KO murine embryonic fibroblast (MEFs) with indicated antibodies. (**E**) Immunoblots of KO MEFs treated for 24h by vehicle and rapamycin with indicated antibodies. (**F**) Immunoblots of the bone marrow cells (BMs), spleen, bone marrow derived mast cells (BMMCs), peritoneal fluid cells (PFCs) and bone marrow derived macrophages (BMDMs) from WT and TSC2 KO mice with indicated antibodies. Data was analyzed by unpaired Student t test. Data are shown as means ± s.e.m. * p<0.05.

### CCL24 expression is suppressed by impaired STAT3 pathway in mTORC1-activated fibroblasts

To elucidate the mechanism driving CCL24 downregulation in TSC1/2-deficient fibroblasts, we conducted transcriptomic profiling of WT and KO MEFs. GSEA revealed that the mTOR pathway and Myc targets were the most enriched gene sets in the KO MEFs and that the IL6-JAK-STAT3 pathway, a major inflammation-regulated pathway, was significantly downregulated (Fig. 5A). Western blot analysis of STAT3 phosphorylation at Y705 demonstrated decreased p-STAT3α and increased p-STAT3β isoforms in TSC2 KO MEFs (Fig. 5B), suggesting dysregulated STAT3 activity. In addition, we found that CCL24 mRNA levels were dramatically reduced in KO MEFs (Fig. 5C), indicating that CCL24 loss might be due to reduced transcriptional activity of hypo-phosphorylated STAT3. To validate our hypothesis, we performed the dual luciferase reporter assay to demonstrate that phosphorylated STAT3 at Y705 is the transcription factor of CCL24. We cloned the CCL24 promoter region which is located within the 4500 base-pairs(bp) upstream of CCL24 transcription start site (TSS) (-4500-0) into the plasmid pGL3.0 and co-transfected with pRL-TK and different forms of Flag-STAT3 (WT, Y705D, Y705A). As indicated, cells expressing STAT3-Y705D mutant which mimics the phosphorylation of STAT3 at Y705 showed a significantly higher relative luminous intensity compared to those expressing STAT3-Y705A mutant which mimics non-phosphorylation of STAT3 at Y705 (Fig. 5D), supporting the notion that expression of CCL24 is positively regulated by phosphorylated STAT3 at Y705 via transcription. To further map the promoter region of CCL24 gene that binds to Y705 phosphorylated STAT3, we performed the chromatin immunoprecipitation quantitative real-time PCR (ChIP-qPCR) and results revealed that pY705-STAT3 specifically binds to a site located at the 601 base-pairs(bp) upstream of the CCL24 TSS (-601) (Fig. 5E and Fig. EV5). These findings demonstrate that mTORC1 activation in fibroblasts leads to the decrease of CCL24 through impaired STAT3 phosphorylation and transcriptional activity, revealing a novel molecular mechanism contributing to sarcoid pathogenesis.

**Figure 5.**
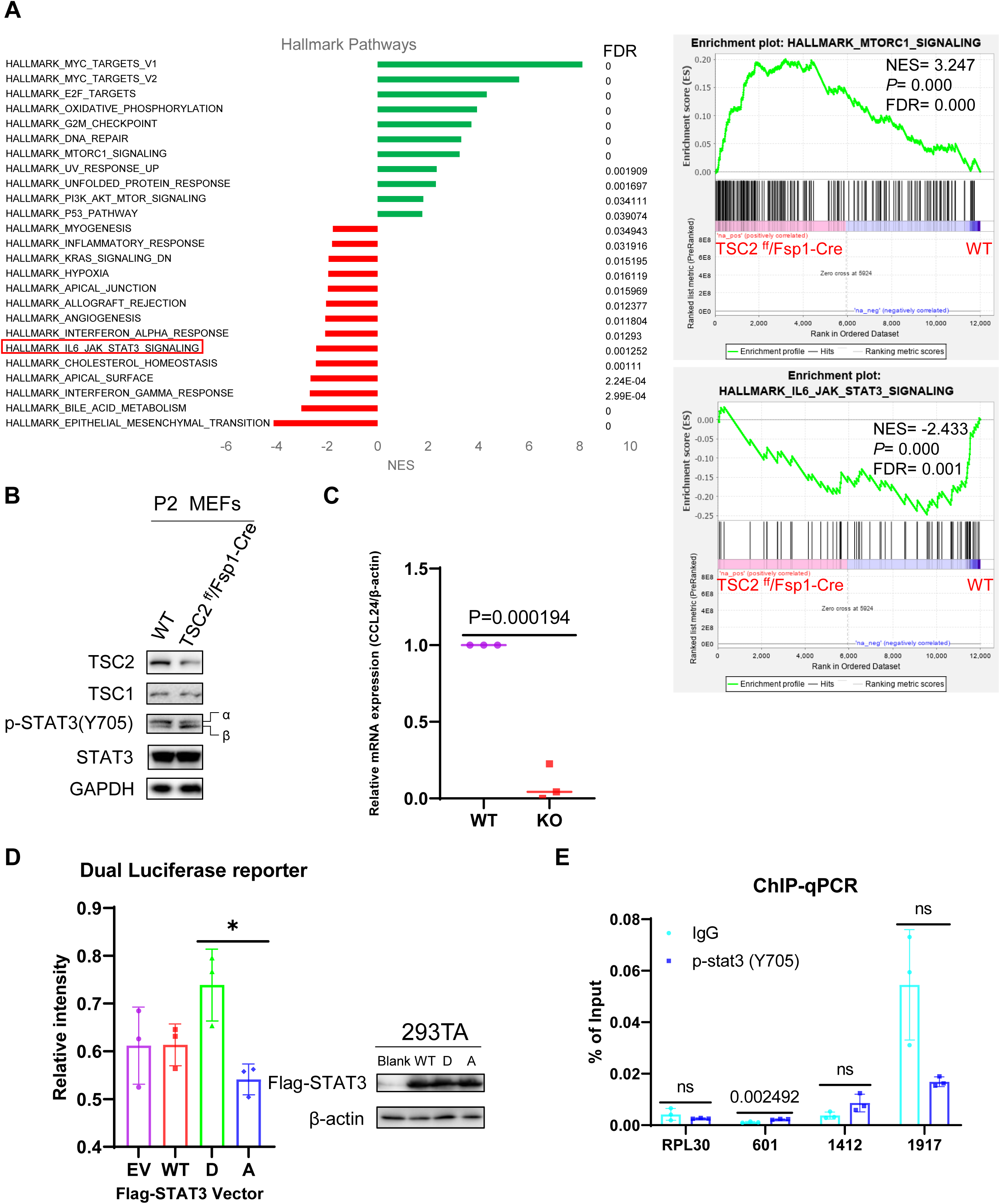
CCL24/eotaxin-2 is regulated by phosphorylated STAT3 (Y705). (**A**) Gene set enrichment analysis (GSEA) comparing WT and KO MEFs, showing indicated normalized enrichment scores (NES) and false discovery rates (FDR). (**B**) Immunoblotting analysis of phosphorylated STAT3 (Y705) levels in WT and TSC2 KO MEFs. (**C**) Relative mRNA expression of CCL24 in WT and TSC2 KO MEFs as measured by qRT-PCR. (**D**) Dual-luciferase reporter assay in HEK293T cells co-transfected with a CCL24 promoter-driven luciferase construct and either empty vector, wild-type (WT) STAT3, phosphomimic STAT3-Y705D, or non-phospho STAT3-Y705A. (**E**) Chromatin immunoprecipitation (ChIP) followed by qPCR analysis showing binding of pY705-STAT3 to the CCL24 promoter region in MEFs. Data was analyzed by unpaired Student t test. Data are shown as means ± s.e.m. * p<0.05.

### CCR3 is upregulated in sarcoid macrophages and contributes to CCL24 loss

We next analyzed CCL24-associated signaling, Biocarta_CCR3 pathway among the immune cells from the lungs and found that CCR3 pathway was significantly upregulated in interstitial macrophages (Int_Macro) from KO lungs (Fig. 6A). These findings led us to hypothesize that increased CCR3 expression in KO interstitial macrophages enhances CCL24 binding, contributing to the marked loss of circulating CCL24 and amplifying its utility as a sensitive biomarker for sarcoidosis. To confirm our hypothesis, we examined CCR3 expression in lung, spleen, liver, thymus, BMDMs and MEFs, and found that CCR3 was significantly upregulated in the lung, spleen, liver, thymus and BMDMs of KO mice, but not in MEFs (Fig. 6B). To explore the clinical relevance, we analyzed gene expression data from the GSE109516 dataset (Vukmirovic *et al*, 2021) to analyze the transcriptional level CCR3 and CCL24 in the BAL and peripheral blood mononuclear cells (PBMCs) from Scadding stage of sarcoid cohort (Fig. EV6A). CCR3 expression was significantly elevated in BAL samples from late-stage sarcoidosis patients (Fig.6C and Fig. EV6B) and showed a marginally significant upward trend in PBMCs across disease progression (Fig. 6D and Fig. EV6C). By contrast, CCL24 mRNA levels remained unchanged in both BAL and PBMCs across sarcoidosis stages compared to healthy controls (Fig. EV6D, E), suggesting the binding to CCR3 may explain its loss. Together, these results suggest that CCL24 loss in sarcoidosis arises from both decreased productions, primarily in fibroblasts and increased expression of CCR3 in sarcoidosis, particularly in sarcoid macrophages (Fig. 6E).

**Figure 6.**
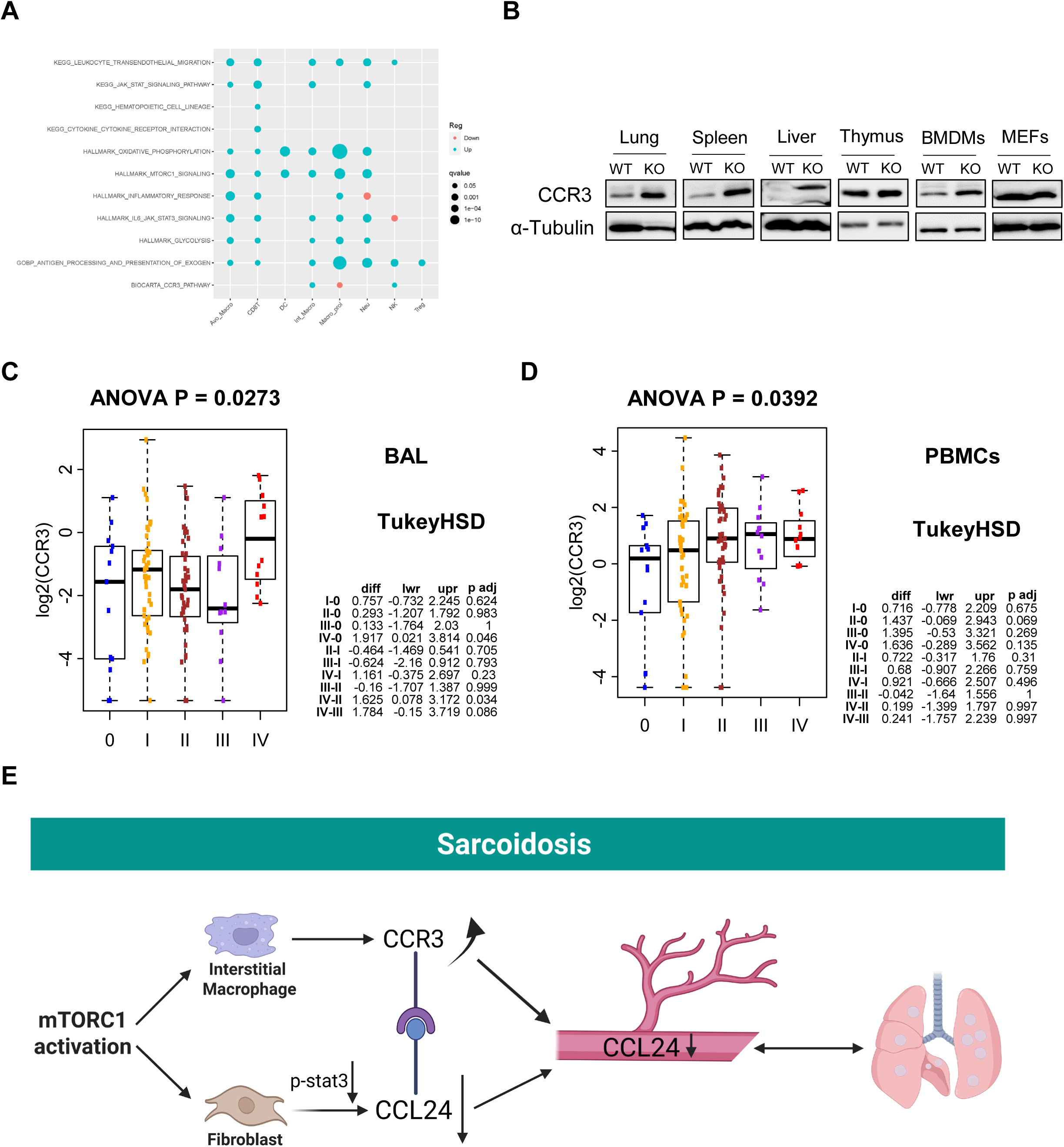
CCR3 is promoted in TSC1/2 ^fl/fl^; Fsp1-cre mice. (**A**) GESA analysis for the immune cells from WT and KO lungs. (**B**) Immunoblotting analysis of CCR3 expression for lung, spleen, liver, thymus, BMDMs and MEFs from WT and KO mice. Box plot together with the corresponding ANOVA p-value to compare the expression of CCR3 among the stages 0–IV within the tissue type of BAL (**C**) and PBMCs (**D**). To adjust for multiple pair-wise comparisons among the stages, the table of statistics of Tukey’s Honestly Significant Difference (HSD) test between each pair of stages is attached to the boxplot. All the comparisons are based on the log2-transformed RNA-Seq expressions (TPM-normalized). Before taking the log2 transformation, the zero expression of CCR3 is imputed by its none-zero minimum expression. (**E**) Working model for the sarcoidosis in TSC1/2 ^fl/fl^; Fsp1-cre mice.

These findings reveal a dual mechanism underlying CCL24 downregulation in sarcoidosis, involving impaired fibroblast-derived secretion and enhanced macrophage receptor-mediated binding, underscoring the functional relevance of the CCL24–CCR3 axis.

### Targeting the CCL24-CCR3 axis attenuates granuloma burden and inflammation in sarcoidosis

Given the critical role of mTOR signaling in granuloma formation and maintenance (Elliott *et al., 2021*; Linke, 2017), we treated TSC1/2 KO mice with the mTORC1 inhibitor rapamycin via intraperitoneal (IP) injection. Rapamycin treatment markedly ameliorated sarcoid-like symptoms, including a reduction in inflammatory signs and promotion of fur regrowth (Fig. 7A). Rapamycin treatment also significantly decreased spleen size (Fig. 7B) and reduced granulomatous infiltration in the lungs (Fig. 7C). Despite these improvements, rapamycin did not restore plasma CCL24 levels at 24 hours post-injection (Fig. 7D). This is likely due to its rapid systemic absorption, as previously reported (Comas *et al*, 2012). Consistent with this, rapamycin significantly increased circulating CCL24 levels at earlier time points, like 2, 4, and 8 hours after administration (Fig. 7E).

**Figure 7.**
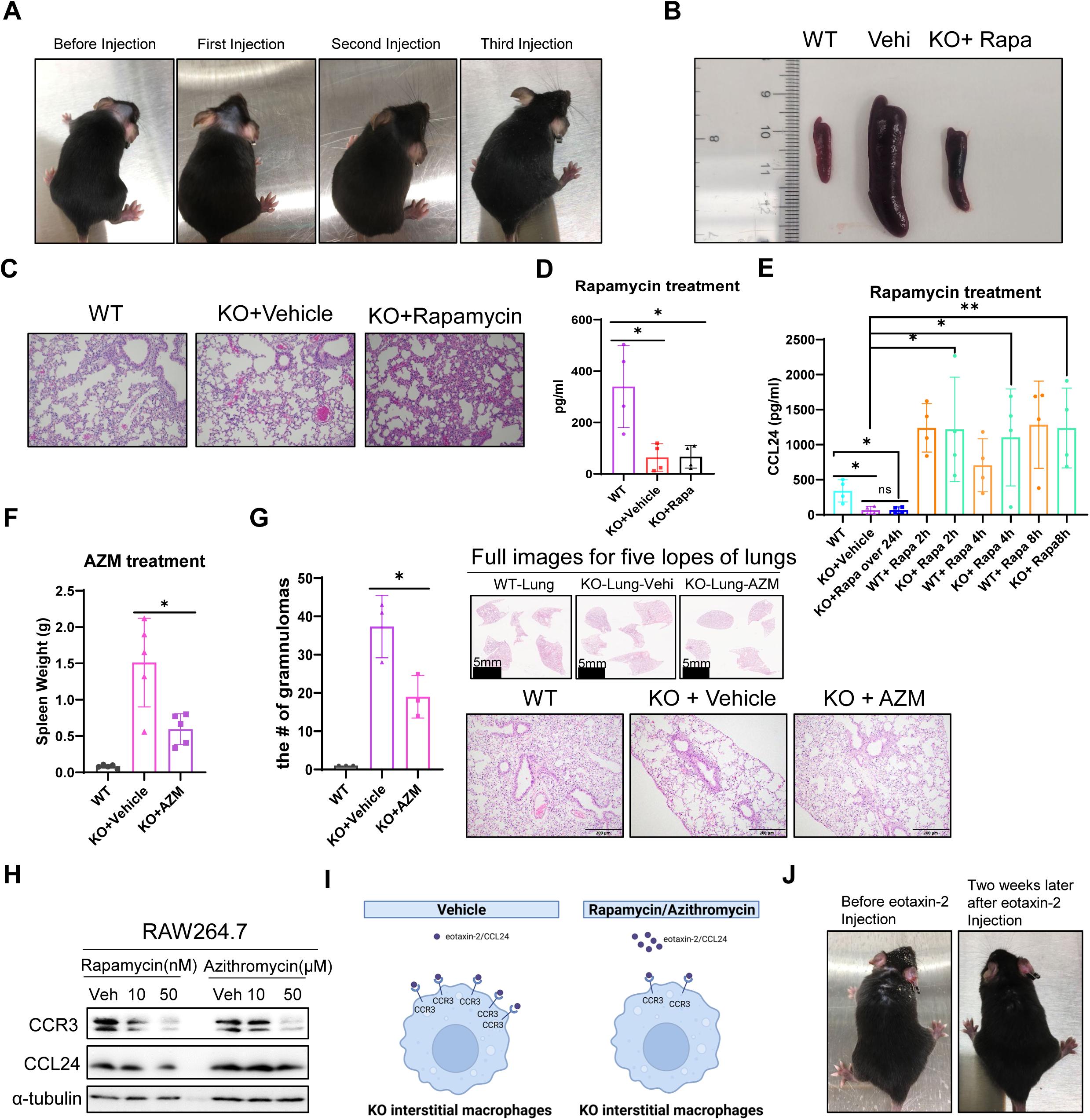
Drug treatment for TSC1/2 ^fl/fl^; Fsp1-cre mice. (**A**) Images of mice injected by rapamycin twice within one week into TSC2 KO mice at the age of 4 months. (**B**) Image of spleens from WT and TSC2 KO mice treated by vehicle (vehi) and rapamycin (rapa). Results are representative of at least 6 mice per genotype. (**C**) H&E staining images of lungs from WT and TSC2 KO mice treated by vehicle and rapamycin. Scale bar,100µm. Results are representative of at least 6 mice per genotype. (**D**) ELISA analysis of CCL24 for the blood plasma from WT and TSC2 KO mice treated by vehicle and rapamycin for over 24 hours. (**E**) ELISA analysis of CCL24 for the blood plasma from WT and TSC2 KO mice treated by vehicle and rapamycin for 2h, 4h and 8h. (**F**) Spleen weights of WT and KO mice treated by azithromycin (AZM) daily for two weeks. (**G**) Granulomas counting for the lungs from WT and KO mice treated by vehicle and AZM. Upper: H&E images of total five lopes for each lung, scale bar=5mm. Lower: H&E images for representative lungs, scale bar=200µm. Results are representative of at least 6 mice per genotype. (**H**) Immunoblotting analysis for the protein levels of CCR3 and CCL24 in RAW264.7 cells upon treatment with different concentrations of rapamycin and azithromycin. (**I**) Schematic of rapamycin/azithromycin treatment for TSC1/2 KO interstitial macrophages to reduce CCR3 and release CCL24. (**J**) The images of TSC2 KO mice before and after injection of recombinant CCL24 for two weeks. Data was analyzed by unpaired Student t test. Data are shown as means ±s.e.m. * p<0.05, ** p<0.01.

To explore additional therapeutic strategies, we investigated azithromycin (AZM), a clinically approved antibiotic known to modulate macrophage polarization from a pro-inflammatory (M1) to anti-inflammatory (M2) phenotype (Haydar *et al*, 2019). AZM administration in KO mice reduced spleen size (Fig.7F) and ameliorated peripheral inflammation, including tail and footpad swelling (Fig. EV7A). Unlike rapamycin, AZM did not promote fur regrowth, suggesting it primarily targets macrophage-driven inflammation rather than mast cell-associated processes involved in hair cycling (Paus *et al*, 1994). AZM treatment also significantly decreased lung granuloma burden (Fig. 7G), although CCL24 levels in serum remained suppressed at 24 hours (Fig.EV7B), likely reflecting similar pharmacokinetic absorption dynamics as rapamycin. To further delineate the mechanism, we treated the RAW264.7 murine macrophage cell line with rapamycin or AZM. While neither drug altered CCL24 expression directly, both significantly reduced CCR3 expression (Fig. 7H). These results suggest that rapamycin and AZM may downregulate CCR3 in macrophages, thereby reducing CCL24 binding and potentially increasing its extracellular availability (Fig. 7I). Given that CCR3 serves as a high-affinity receptor for CCL24, its suppression may relieve the cytokine sink effect, indirectly enhancing CCL24-mediated signaling. While AZM shows therapeutic potential in modulating the CCL24-CCR3 axis, prolonged or higher-dose regimens may be necessary to fully restore CCL24 signaling in sarcoidosis.

To further confirm CCL24 loss in KO mice resulting in the maintenance and progression of pulmonary sarcoid phenotypes, we neutralized CCL24 by injecting CCL24 antibodies into WT mice. Within 48 hours, these mice exhibited white blood cell profiles resembling those of KO animals (Fig. EV7C), supporting the functional significance of CCL24 depletion in promoting systemic inflammation. Conversely, administration of recombinant CCL24 to KO mice effectively reduced inflammatory phenotypes (Fig.7J), directly linking CCL24 deficiency to sarcoid-like disease manifestations. Altogether, these results establish that therapeutic modulation of the CCL24-CCR3 axis, through mTORC1 inhibition, macrophage targeting, or recombinant cytokine supplementation, represents a promising strategy to mitigate granuloma burden and inflammation in sarcoidosis.

## Discussion

In this study, we established and characterized a novel murine model of sarcoidosis by deleting TSC1 or TSC2 using a fibroblast-specific Fsp1-Cre driver. Notably, Fsp1-Cre activity was not restricted to fibroblasts but also extended to hematopoietic lineages, enabling a more physiologically relevant model of sarcoidosis compared to previous Lyz2-Cre models limited to macrophages (Linke *et al*., 2017). In addition, the polarized macrophages were not markedly elevated in TSC2 ^fl/fl^;Lyz-cre BMDMs (Linke *et al*., 2017), but significantly increased in our TSC2 ^fl/fl^;Fsp1-cre BM (Fig. EV2A). This broader expression pattern recapitulates the complex multicellular interactions involved in granuloma formation and maintenance, reflecting more accurately the pathophysiology observed in human sarcoidosis (Krausgruber *et al*, 2023a).

A central and clinically translatable discovery of our study is the consistent and significant downregulation of CCL24 (eotaxin-2) in both the TSC1/2 KO mouse model and sera from human sarcoidosis patients. While genetic predisposition for CCL24 downregulation in sarcoidosis has been suggested (Meguro *et al*., 2020), the molecular and cellular mechanisms underlying this phenomenon remained unexplored. Our study fills this gap by identifying a mechanistic link between mTORC1 activation in fibroblasts, impaired STAT3 signaling, and transcriptional suppression of CCL24.

Importantly, our model faithfully recapitulates multiple clinical manifestations of sarcoidosis, including pulmonary granulomas, splenomegaly (Kawano *et al*, 2012; Kyriakos Saad *et al*, 2024; Saito *et al*, 2020), hepatic infiltration (Giovinale *et al*, 2009; Kikuchi *et al*, 2024), skin involvement, and hematopoietic abnormalities (Fraser *et al*, 2024; Hiranuma *et al*, 2024). We also demonstrate that granuloma formation is driven not only by hyperactive fibroblasts but also by interstitial and alveolar macrophages and dendritic cells with mTORC1-dependent transcriptional reprogramming. These immune cells exhibit upregulated CCR3 expression, the receptor for CCL24, suggesting a feed-forward loop in which elevated CCR3 enhances chemokine binding, further depleting circulating CCL24 and sustaining granulomatous inflammation.

Therapeutically, we demonstrate that targeting the CCL24-CCR3 axis can attenuate disease progression. Treatment with rapamycin, a clinically available mTORC1 inhibitor, or azithromycin, an FDA-approved macrolide antibiotic with immunomodulatory properties, significantly reduced granuloma burden and inflammation. Both drugs suppressed CCR3 expression in macrophages, potentially increasing CCL24 releasing. While neither fully restored CCL24 levels with single-dose administration, temporal analysis revealed transient restoration with rapamycin, suggesting pharmacokinetic factors and duration of treatment as key considerations. Furthermore, direct neutralization of CCL24 in wild-type mice recapitulated sarcoidosis-like immune profiles, while recombinant CCL24 administration reversed disease features in KO mice, establishing a causal role for CCL24 depletion in granuloma maintenance. Specifically, CCL24 antibody injection markedly increased the percentage of monocytes in peripheral blood from WT mice (Fig. EV7C), suggesting the potential role of CCL24 in regulating the homeostasis of monocytes, which needs to be further investigated.

These findings have major implications for sarcoidosis treatment. Current therapeutic regimens rely heavily on broad immunosuppression, primarily corticosteroids, which are associated with significant adverse effects and limited long-term efficacy (Megari, 2013). Our study underscores the potential for pathway-specific, immunomodulatory interventions that target the CCL24-CCR3 axis or mTORC1 signaling. The success of rapamycin in a reported sarcoidosis case (Manzia, 2011) and the repositioning potential of azithromycin further support this direction. These data suggest the need to potentially explore these combination therapies in patients with sarcoidosis.

Finally, our research bridges basic mechanistic insights with clinical application. We validated key findings from our murine model in patient sera, confirming the translational relevance of CCL24 as both a diagnostic biomarker and a therapeutic target. By elucidating the cellular and molecular underpinnings of granuloma formation and persistence, and by establishing a robust preclinical platform, this study lays the groundwork for rationally designed therapies and precision medicine approaches in sarcoidosis.

## Methods

### Mouse strains

TSC1^flox/flox^ mice (STOCK Tsc1tm1Djk/J, strain #005680), TSC2 ^flox/flox^ mice (STOCK Tsc2tm1.1Mjg/J, strain # 027458), Fsp1-Cre mice (BALB/c-Tg(S100a4-cre)1Egn/YunkJ, strain # 012641), mTmG mice (STOCK Gt (ROSA)26Sortm4(ACTB-tdTomato, -EGFP) Luo/J, strain#:007576) were purchased from Jackson Laboratory. TSC1/2 ^fl/fl^ mice were crossed with Fsp1-cre/+ mice and the pups were backcrossed to obtain TSC1/2^fl/fl^/Fsp1-cre/+ (denoted TSC1/2 KO) or TSC1/2^fl/fl^, Fsp1-Cre/+, TSC1/2^fl/+^/Fsp1-cre (denoted WT) littermates. Male and female mice were used randomly, and no major sex-specific differences were observed. Groups in individual experiments were sex-matched and age-matched. The ear or tail pieces were harvested for the genotyping according to Jackson Laboratory’s instructions. All mouse studies were approved by the official Kentucky ethics committee for animal experiments and University of Kentucky (UKy) Animal Care and Use Committee (Protocol no: 2020–3685).

### Human sample collection

All human subjects were recruited under protocols approved by the University of Kentucky (UKy) Institutional Review Board (IRB#51510). Written informed consent was obtained from all participants in accordance with the Declaration of Helsinki. Sarcoidosis diagnoses were confirmed by pathologists at the University of Kentucky Medical Center. Blood samples were collected through the UKy Center for Clinical and Translational Science (CCTS). This study included blood samples from 21 healthy individuals and 9 treatment-naïve patients with sarcoidosis.

### Cell culture

Murine macrophage RAW264.7 cells, 293T cells were purchased from ATCC and cultured according to the ATCC’s handling information. The WT and TSC1 or TSC2 KO MEFs isolation was performed according to the method described by Xu (Xu, 2005). Briefly, mouse breeder pairs were set up and females were checked for copulatory plugs the following morning. On day 13.5, the pregnant female mouse was euthanized via cervical dislocation, and the intact uterus was extracted and cut into sections between each embryo. Then each embryo was separated from the placenta, membranes, and umbilical cord. The bulk of the CNS tissue was removed by severing the head above the level of the oral cavity and saved for genotyping. Forceps were used to remove the dark red tissue such that most remaining cells were fibroblasts. In a new petri dish, the embryo body was minced and then transferred to a 15 ml conical tube with 5 ml culture media and centrifuged. The supernatant was aspirated and the tissue washed with 1 × PBS. The pellet was re-suspended into 2 ml MEFs culture medium (DMEM with 10% FBS, 1% P/S). The embryonic cell and media mixture were then transferred to tissue culture dishes with fresh culture media and placed in 37°C incubator to grow. All the cells were cultured in the incubator at 37 °C with humidified 5% CO2. Bone marrow cells from 4-6-month-old WT and KO littermates were flushed from femurs, and tibias and cultured in the 10cm dishes with 10mL following media: DMEM high glucose (Sigma), 10% low endotoxin FBS (Gibco), 2 mM L-glutamine (Sigma), 100 U/ml penicillin (Sigma), 100 µg/ml streptomycin (Sigma), 50 µg/ml β-mercaptothion (Gibco). Floating cells were removed, and adherent cells were split 1:2 in fresh DMEM media. Differentiated BMDM (more than 90% of the cells were positive for F4/80 and CD11b) were seeded in DMEM supplemented with 25ng/ml CSF-1 (PeproTech) and 5.5µM ᵦ-mercaptothion for 4 weeks. For c-Kit^+^ and FcγRII/RIII^+^(CD16/32^+^) mast cell differentiation, bone marrow cells were cultured for 4 weeks on cell culture-treated dishes (one 6-well plate per mouse and 4 ml per well) in media: RPMI 1640 (Sigma), 10% low endotoxin FBS (Gibco), 2 mM L-glutamine (Sigma), 100 U/ml penicillin (Sigma), 100 µg/ml streptomycin (Sigma), 50 µg/ml β-mercaptothion (Gibco) supplemented with 5 ng/ml IL-3 (PeproTech) and 50ng/ml SCF(PeproTech). The media was refreshed every 48 hours.

### Collection Procedure

The peripheral blood (PB) of the mice was collected through the tail vein. The blood cells were counted using Hemavet 950 (Drew Scientific) or ProCyte Dx Hematology Analyzer (IDEXX).

Mouse bronchoalveolar lavage (BAL) fluid was collected according to a published protocol (Au - Van Hoecke *et al*, 2017). Briefly, the mice were euthanized by isoflurane and placed on their backs on a surgical plate, fixed by pinning down the limbs. The neck was sprayed with 70% ethanol and the trachea was exposed by a scalpel and then carefully punctured. The BAL were collected by a 1 ml syringe with 19 g blunt needle after injection of 1 ml ice-cold PBS into the trachea.

Mouse peritoneal fluids (PFL) were collected according to a published protocol (Au - Ray & Au - Dittel, 2010). Briefly, the mice were euthanized and lay down on the styrofoam block on their backs making the block at approximately a 30-degree angle with the head faced up. The outer skin of the peritoneum was cut after being sprayed with 70% ethanol, and then 5 ml ice-cold PBS with 3% fetal calf serum (FCS) was injected into the peritoneal cavity using a 27g needle. The peritoneal fluids were collected using a 5 ml syringe with 25 g needle after the peritoneum was gently massaged to dislodge any attached cells into the PBS solution.

### Flow cytometry and cell sorting

Lungs, livers and spleens were chopped into small pieces and ground with the plastic tissue grinder. The ground spleens, peripheral blood cells and flushed bone marrow cells were suspended by the red blood lysis buffer (NH4Cl 0.15 M, KHCO3 10 mM, EDTA 0.1 mM). The tissue cells were collected through a 40-µm cell strainer in cold PBS or PBS containing 2% fetal bovine sera (FBS). Antibodies from BioLegend were CD45-APC (#103112), CD45.2-Brilliant Violet 510(BV510) (#109837), F4/80-Alexa Flour 488 (#123120), F4/80-BV605 (#123133), CD11b (Mac1)-FITC (#101206), CD11b (Mac1)-APC (#101212), Ly-6G/Ly-6C(Gr-1)-FITC (#108406), Ly-6C-BV711 (#128037), CD11c -FITC (#117306), CD3-PE (#100206), CD3-FITC (#100204), CD3-PECy7 (#100220), Ly-6A/E(Sca-1)-APC (#122511), CD117(c-Kit)-BV650 (#105853), B220-PE (#103208), CD4-APC (#116014), CD4-APCCy7 (#100526), CD8a-FITC (#100706), CD8a-PECy7 (#100722), CD16/32-FITC (#101306), CD115-APC (#135510), CD206-APC (#141707), CD68-APC (#137008), Siglec F(CD170)-bv421 (#155509). I-A/I-E (MHC class ‖)-FITC (# 562009), I-A/I-E (MHC class ‖)-PE (# 562010) were obtained from BD Bioscience. CD45-PE (#12-0451-82), CD3e-FITC (#11-0031-82), CD48-FITC (#11-0481-82), CD150-APC (#17-1501-81), CD117(c-Kit)-APCCy7 (#47-1171-82), L6-6A/E(Sca-1)-PECy7 (#25-5981-82), Ter-119-PE (#12-5921-82), CD11b(Mac-1)-PE (#12-0112-83), Ly6G/Ly-6C(Gr-1)-PE (#12-5931-83), CD34-Alexa Fluor 488 (# 53-0341-82) were obtained from eBioscience(ThermoFisher). All the conjugated antibodies were diluted at the ratio of 1:100∼1:200 for flow cytometry. Data was obtained on Symphony A3 (BD Biosciences) or CytoFLEX LX (Beckman Coulter) and processed using FlowJo (Tree Star Inc, Ashland, OR). Gating for eosinophils sub-populations in the tissues was performed as previously described (Joanne *et al*., 2014). T-cells were defined as CD45^+^CD3e^+^ and positive for CD4 or CD8 respectively, B-cells were defined as CD45^+^B220^+^, myeloid cells were defined as Mac1^+^Gr1^+^, polarized macrophages were defined as Mac1^+^F4/80^+^, mast cells were defined as c-Kit^+^ CD16/32^+^.The lung dissociated cells were kept in RPMI 1640 containing 10% FBS, stained with 7-Aminoactinomycin D (7-AAD) and then loaded on SY3200 Cell Sorter (Sony Biotechnology) to sort about 200,000 cells for the single-cell RNA sequencing (scRNA-seq).

### Western Blot

The tissues were lyophilized by liquid nitrogen, ground by the tissue grinders or a porcelain mortar with pestle. The cells were washed and scraped in ice-cold PBS. The pellet of tissues or cells were dissolved in lysis buffer (EMD Millipore) supplemented with protease and phosphatase inhibitors (Roche) for 10 min on ice and then sonicated at 40% power. Protein concentration was measured by Pierce BCA Protein Assay kit (ThermoFisher) and equal amounts (20∼50 µg) of denaturized lysate were resolved on 7.5–12% SDS-PAGE and transferred to nitrocellulose (NC) or polyvinylidene difluoride (PVDF) membranes. Membranes were blocked in 5% low-fat milk for 1 h at room temperature (RT) and incubated with primary antibodies at 4 °C overnight. HRP-conjugated secondary antibodies (CST) 1:5,000 were applied for 1 h at RT in 5% low-fat milk and proteins were visualized using ECL substrate (ThermoFisher).

Primary antibodies were TSC1(D43E2), TSC2 (D93F12), p-Akt S473 (E4U3U), p-S6 S240/244 (D68F8), S6 (54D2), p-Akt T308 (D25E6), p-AKT T450 (D5G4), pan-Akt (40D4), β-tubulin (9F3), p-TSC2 T1462 (5B12), STAT3 (124H6), p-STAT3 (Y705) (D3A7), mTOR (7C10), p-mTOR (S2481), β-actin (8H10D10) obtained from Cell Signaling Technology and diluted at 1:1000. Monoclonal ANTI-FLAG M2 antibodies were purchased from Millipore Sigma and diluted at 1:3000. CCL24 were obtained from Proteintech (Cat#: 22306-1-AP), diluted at 1:200 and Novus (Cat#: NBP2-12109), diluted at 1:100.

### Immunohistochemistry (IHC)

The tissues were collected freshly and fixed with 4% formaldehyde overnight. Then, the fixed tissues were delivered to Biospecimen Procurement & Translational Pathology Shared Resource Facility (BPTP SRF) for paraffin embedding, cutting and sectioning using a cryostat (Leica) onto Superfrost Plus Microscope Slides (Fisherbrand), following Hematoxylin and Eosin (H&E) staining. VECTASTAIN Elite ABC Universal PLUS Kit was purchased from VECTOR laboratories and IHC was performed according to the manufacturer’s instructions using the following primary antibodies as well as the indicated dilutions to stain slides overnight at 4 °C: p-S6 (S 240/244) (D68F8, Cell Signaling Technology). Immunostaining experiments were performed on at least three mice with the same genotyping and images were acquired and identified by the Nikon ECLIPSE Ti2 microscope with at least 10 fields of view in each.

### mRNA expression analysis

RNA was isolated by the RNeasy Mini Kit (QIAGEN) according to the manufacturer’s instructions. Equal amounts of RNA were transcribed to cDNA using QuantiTect Rev. Transcription Kit (QIAGEN). mRNA levels were determined by FastStart SYBR Green Master (Roche) on a StepOnePlus or QuantStudio 3 Real-Time PCR System. Relative expressions were normalized to β-actin or GAPDH.

### The enzyme-linked immunosorbent assay (ELISA)

Mouse CCL24 ELISA Kit (Cat#: EMCCL24) and Human CCL24 ELISA Kit (Cat#: EHCCL24) were purchased from ThermoFisher. The analysis of human or mouse CCL24 were followed by the manufacturer’s instructions.

### Inflammatory Chemokine Array

Mouse Inflammation Array G1 and Mouse Inflammation Array G1 kits were purchased from RayBiotech. The inflammatory cytokines of mouse blood plasma and bone marrow plasma were assayed by the manufacturer’s instructions.

### RNA-Seq data analysis

Sequencing reads were trimmed and filtered using Trimmomatic (V0.39) to remove adapters and low-quality reads (Bolger *et al*, 2014). Reads were mapped to Ensembl GRCm38 (release 100) transcripts annotation using RSEM (Li & Dewey, 2011). RSEM results normalization and differential expression analyses were performed using the R package edgeR (Robinson *et al*, 2009). Significantly up/downregulated genes were determined as fold change >= 2 and q-value < 0.05. Gene set enrichment analysis was performed using GSEA software and the Hallmark gene sets in the Molecular Signature Database (MSigDB) (Subramanian, 2005).

### Dual luciferase reporter assay

CCL24 promoter region (-4500-0) was cloned into the plasmid pGL3.0 (Promega) based on the extracted cDNA from RAW264.7 cells, HT29 cells and H116 cells. Flag-STAT3 was purchased from SinoBiological and Y705A, and Y705D were mutated using the New England Biolabs (NEB) Q5 Site-Directed mutagenesis Kit. The dual luciferase assay was performed by sequentially measuring the firefly and Renilla luciferase activities of the same sample after transfection of CCL24 promoter and empty vector, WT STAT3, D mutation, A mutation, respectively in 293T cells, with the results expressed as the ratio of firefly to Renilla luciferase activity (Fluc/Rluc). Primers used are:

hCCL24-promoter-F: CG ACGCGT GGGTTGGTAATCCCTGCCTT

hCCL24-promoter-R: CCG CTCGAG GTCTCAGAGAGCAGAAGCAC

STAT3Y705D-F: CGCTGCCCCAgacCTGAAGACCA

STAT3Y705D-R: CTACCTGGGTCAGCTTCAGGATG

STAT3Y705A-F: CGCTGCCCCAgccCTGAAGACCA

STAT3Y705A-R: CTACCTGGGTCAGCTTCAG

ChIP, ChIP-PCR and ChIP-qPCR

ChIP, ChIP-PCR, and ChIP-qPCR were performed following the manufacturer’s instructions (CST,9003S) and the previously described protocol (Fong *et al*, 2022). 1×10^7^ RAW264.7 cells were crosslinked with 1% formaldehyde in fixation buffer (50mM HEPES pH7.5, 100mM NaCl, 0.5mM EDTA) for 10 min at room temperature with gentle rotation and then quenched with 0.125 M glycine for 5 min. Nuclei was digested by micrococcal nuclease (CST, Cat#10011), and the supernatant was used for immunoprecipitation with the indicated antibody. The primers which were designed based on JASPAR and Primer 3 are:

CCL24-601-F: CTGGGAACAAGCCATCATCT

CCL24-601-R: CACTCTACCTGCCCATCCAT

CCL24-1412-F: AACATTTTCCCTTGGCTGTG

CCL24-1412-R: CTTGTCCAGTGGAGTCAGCA

CCL24-1917-F: GGCTCTGTGTCTGCAGTTGA

CCL24-1917-R: AGATGAGGGGATGGTCACAG

### Treatment

Rapamycin (everolimus) was dissolved in pure ethanol to a stock concentration of 40 mg/ml and stored at -20°C. The stock rapamycin was further diluted by vehicle to the final concentration of 1 mg/ml before use. 8 to 20-week-old TSC1/2^fl/fl^, Fsp1-Cre mice were injected with 5 mg/kg body weight everolimus or provided placebo daily for three weeks. Different mice were bled after treatment with rapamycin at 2h, 4h, 8h and 24h and analyzed by flow cytometry and CCL24 ELISA or sIL2R ELISA. The vehicle was prepared with sterile water containing 0.25% Tween 80 and 0.25% PEG400.

Azithromycin was ordered from EPICpharma by the Division of Laboratory Animal Resources (DLAR) of the University of Kentucky (UKy) and dissolved in sterile water to the final suspension of 100 mg per 5 mL. The mouse was administered 150 µL azithromycin suspension daily through gavage and monitored twice a week. Different mice were bled at 2h, 4 h, 8h and 24h for analysis by flow cytometry and CCL24 ELISA.

Recombinant mouse CCL24/eotaxin-2 were purchased from BioLegend (Cat# 585106) and dissolved in sterile water to a final concentration of 100 ng/µL. CCL24 was administered to mice daily via intraperitoneal (IP) injection with 100 µl/mouse as previously reported (Menzies-Gow *et al*, 2002). The mice were monitored twice a week and bled for analysis after injection for 2 days or 2 weeks by flow cytometry.

### Mouse lung dissection, dissociation and single cell preparation

The mice were anesthetized with an endpoint dose of ketamine/xylazine (25G needle on 1mL syringe, IP injection) and then affixed to a dissecting platform with pins and sprayed with 70% ethanol after ensuring lack of sensation/reflex. The stomach skin was opened to expose the base of the sternum, and the ribcage was carefully cut up to the neck by holding the base of the sternum with forceps. Then, the skin was cut from the center line, and the two sides of the rib cage were folded to open and secured with pins. The mice were additionally euthanized by removing the heart and then exposing the trachea by placing forceps under the trachea. To dissect out the trachea and lungs, gently tug on the trachea while snipping away the connective tissue; leave the lungs and trachea intact and place in 12 well dishes on ice with Ca^2+^/Mg^2+^-free PBS. The lungs were then moved to a 1.5mL tube and minced using surgical scissors to a homogenous paste, suspended by 1mL Ca^2+^/Mg^2+^-free PBS using 1ml pipette tip with a wide bore and moved to a 50cc conical tube for further dissociation. Using a 5mL stripette with 4mL additional Ca^2+^/Mg^2+^-free PBS and triturated aggressively about 10-20 times. We then passed the whole solution through 70µm and 50cc conical filters, flushing with an additional 7mL Ca^2+^/Mg^2+^-free PBS. The cells were centrifuged at 1000rpm in a typical tabletop centrifuge and underwent Red Blood Cell Lysis if necessary. If needed, remove supernatant carefully and resuspend the cell pellet in ∼200 µL of Red Blood Cell lysing solution (0.15 M NH4Cl, 10mM KHCO3, 0.1 mM EDTA, in 1L distilled H2O; filtered with 0.45 µm filter and stored at RT) to lyse 120 sec at RT, adding 1 mL of PF10 and mixing to end lysis, vertexing 20 seconds before pulse spinning to pellet. The supernatant was carefully removed, and the pellet was resuspended in 300µL Ca^2+^/Mg^2+^-free PBS with 10% FBS (PF10). We selected immune cells by adding EpCAM and CD31 antibodies and let them bind for 10 minutes at RT. The cells were then washed with 1mL PF10, pulse spanned and resuspended in 80µL PF10 and 20µL anti-rat IgG beads. After incubating 15min at 4 °C, the cells were washed by adding 1mL of PF10, following a brief vortex and pulse spin. We then carefully aspirated liquid, resuspended the cells in 500µL PF10 and flushed the magnet column which was placed in a 15cc tube with 3mL PF10. We collected a total of 9 ml PF10 by repeating this process twice. After centrifuging at 1200 rpm for 5 min at 4°C, we then resuspended the pellet in 200-600 µL standard media (RPMI + 10% FBS and glutamax, pen/strep) containing 7-AAD to keep viability according to the pellet size. Finally, the single-cell suspensions were placed on ice and delivered to the UKy Flow and scRNA facility.

### Single Cell mRNA Sequencing (scRNA-seq)

Flow sorted single cell suspensions were centrifuged and resuspended in PBS + 10%FBS to target a concentration of approximately 1000 cells/microliter and evaluated using acridine orange and ethidium homodimer to mark cells. Viabilities ranged from 84-92%. Cells were loaded to capture approximately 10,000 cells per sample and partitioned using the 10x Genomics Chromium Controller with the Chromium Next GEM Single Cell 3’ Kit v3.1 (10x Genomics, Pleasanton, CA). cDNA libraries were generated following the manufacturer’s instructions. Libraries were sequenced on an Illumina NovaSeq 6000 (Novogene Corp.) at a read depth of approximately 35,000 read pairs per cell. Sequence data were demultiplexed and processed using Cell Ranger v6.1.1 (10x Genomics). For scRNA-seq, Cell Ranger was used for sequencing read alignment, filtering, barcode counting and unique molecular identifier counting. Seurat was used for data normalization, dimension reduction, clustering and visualization (Butler *et al*, 2018; Stuart *et al*, 2019). An integrated analysis was performed across all samples to identify cell types and quantify their fractions in each sample. The two-sample binomial test was used to compare the fraction of each cell type between Knockout and wildtype samples. MAST was used to identify differentially expressed genes between experimental conditions for each cell type (Finak G, 2015). The gene set enrichment analysis was performed using GSEA based on the ranked gene list obtained the differential expression analysis. Multiple comparisons adjustment was addressed using the Benjamini-Hochberg procedure to control the false discovery rate (FDR).

### scRNA-seq analysis

scRNA-seq gene expression libraries were mapped to mm10 mouse reference (10X Genomics pre-built reference, mm10-2020-A) using Cell Ranger (v6.1.1, 10x Genomics). For each snRNA-seq library, Cell Ranger filtered matrix was subjected to QC filters to remove low quality cells with ≤ 500 features or percentage of mitochondrial transcripts ≥ 10. DoubletFinder (McGinnis *et al*, 2019) analysis was performed on each library separately to identify and filter potential doublets with default parameters. Additionally, cells with features > 7500 or counts > 40,000 were removed. Pass QC cells from the KO and WT groups were combined and processed using the Seurat (v) R package (Stuart T, 2019). Count matrices were log-normalized and scaled. The top 2,000 highly variable genes (HVGs) were selected to define 20 principal components (PCs). Batch integration was performed via Harmony algorithm (v0.1.0) (Korsunsky *et al*, 2019) to the combined data for batch effect corrections with default settings. Neighbor analysis was performed by FindNeighbors function using the first 20 PCs from the Harmony dimension. Clustering was identified with FindClusters function at resolution = 0.8, resulting in 20 clusters. UMAP was calculated based on the Harmony dimension for clusters visualization. Identification of cluster markers was performed using a Wilcoxon rank-sum test by comparing each cluster with the rest of the cells. For differential gene expression analysis for each cell population, we used DESeq2 test (Love *et al*, 2014) in Seurat, to identify the up/down-regulated sets of genes between treatment groups in the cell populations. The two-sample binomial test was used to compare the distribution of cell clusters between the 2 groups. Cell–cell interaction analysis was performed using CellChat (Jin *et al*, 2021).

### Statistical Analysis

Data was analyzed by GraphPad Prism (Version 9) or Microsoft Excel. Data were modified by Photoshop 2024 or Microsoft PowerPoint. The significance of data sets was analyzed using the unpaired two-tailed t-test and p<0.05 was considered significant. NS, is not significant. In comparing gene expressions among the stages of patients, p-values were obtained by ANOVA (analysis of variance).

## Acknowledgments

The research was generously supported by NIH R01 CA157429 (X. Liu), R01 CA196634 (X. Liu), R01 CA264652 (X. Liu), R01 CA256893 (X. Liu), R01 CA266579 (Z. Li). This research was also supported by Biospecimen Procurement & Translational Pathology, Biostatistics and Bioinformatics, Redox Metabolism, and Flow Cytometry and Immune Monitoring Shared Resources of the University of Kentucky Markey Cancer Center (P30CA177558). The research was also supported by the UK IRC grant (#1013171125, P. Sen).

## Author contributions

**Xiongjian Rao**: Conceptualization; Project designation; Investigation; Methodology; Data curation; Writing-original draft; Writing-review and editing. **Jinpeng Liu**: Methodology; Investigation; Bioinformatic analysis. **Derek B Allison**: Visualization; Validation; Methodology. **Douglas A Harrison**: Methodology; Investigation; Resources. **Ka Wing Fong**: Methodology; Resources; Formal analysis. **Yuanyuan Wu & Peng Jia**: Methodology. **Daheng He**: Methodology; Formal analysis. **Zhiguo Li**: Resources; Funding. **Chi Wang**: Methodology; Formal analysis. **Jamie L. Sturgill**: Methodology; Resources; Writing-review and editing. **Parijat Sen**: Validation; Funding; Writing-review and editing. **Xiaoqi Liu**: Supervision; Resources; Funding; Writing-review and editing.

## Disclosure and competing interests

Authors declare that they have no competing interests.

## Data and materials availability

All data needed to evaluate the conclusions in the paper are present in the paper and/or the Supplementary Materials. For more information about the data and reagents listed in this manuscript, please request the corresponding authors. Gene Expression Omnibus Microarray data have been deposited, and the accession code will be released. We used the data from GSE109516 (Vukmirovic *et al*., 2021) to analyze the BAL and PBMCs.

## Expanded View Figure legends

**Figure EV1. Phenotypic characterization of TSC1/2fl/fl; Fsp1-cre (TSC1/2 KO) mice.**

(A) Representative images of TSC1 KO mice displaying inflammation around the ear and neck, as well as swelling of the tail and footpads. (B) Representative images of TSC2 KO mice showing similar inflammatory features in the ear, neck, tail, and footpads. (C) Images comparing WT and TSC2 KO mice demonstrating ocular alopecia. (D) Quantification of spleen weights in WT, TSC1 KO, and TSC2 KO mice. (E) Kaplan–Meier survival curves of WT, TSC1 KO, and TSC2 KO mice. Mice euthanized at humane endpoints were included in the survival analysis. Data was analyzed by unpaired Student t test and are shown as means ± s.e.m. ** p<0.01, **** p<0.0001.

**Figure EV2. TSC1/2 KO mice exhibit systemic immune alterations consistent with sarcoid disease.**

**(A)** Flow cytometric quantification of polarized macrophages in the peripheral blood, bone marrow, spleen, liver, and lung of WT and TSC1/2 KO mice. (**B**) Proportion of mast cells in peritoneal fluid cells from WT and TSC1/2 KO mice. (**C**) Proportion of eosinophils in the liver of WT and TSC1/2 KO mice. (**D**) Proportion of eosinophils in the lung of WT and TSC1/2 KO mice. Data was analyzed by unpaired Student t test. Data are shown as means ± s.e.m. * p<0.05, ** p<0.01, *** p<0.001.

**Figure EV3. Comparison of ligand receptor mapping of WT and TSC KO lung immune cells.**

**(A)** Analysis of the cell interactions by the number and strength for the WT and TSC KO lungs. (**B**) Analysis of the cell interactions of WT and TSC KO lungs by immune cell types. (**C**) The incoming and outgoing interaction strength of WT and TSC KO immune cells from lungs. (**D**) The clusters of the cytokines from the WT and TSC KO lung immune cells. (E- H) The signaling changes of neutrophils (**E**), interstitial macrophages (**F**), CD4+ T cells (**G**), and alveolar macrophages (**H**). (**I**) The overall signaling patterns of the WT and KO lung immune cells. (**J**) The CCL signaling of WT and KO major lung immune cells.

**Figure EV4. CCL24/eotaxin-2 decreases in KO mice.**

**(A)** Quantification for the inflammatory cytokines/chemokines of the mouse blood plasma. (**B**) Quantification for the inflammatory cytokines/chemokines of the mouse bone marrow plasma.

**Figure EV5. CCL24/eotaxin-2 is regulated by Y705 phosphorylated STAT3.**

ChIP-PCR for the lysates of IMR90 cells by using p-STAT3 (Y705) antibodies and the indicated CCL24 promoter primers.

**Figure EV6. The expression of CCR3 and CCL24 among BAL and PBMCs from treatment naïve cohort.**

(**A**) The analyzed sample sizes of treatment naïve stages 0 – IV cohort. (**B**) The 95% confidence intervals for the expression of CCR3 among the stages 0 – IV within the tissue type of BAL. (**C**) The 95% confidence intervals for the expression of CCR3 among the stages 0 – IV within the tissue type of PBMCs. Box plot together with the corresponding ANOVA p-value to compare the expression of CCL24 among the stages 0–IV within the tissue type of BAL (**D**) and PBMCs (**E**). To adjust for multiple pair-wise comparisons among the stages, the table of statistics of Tukey’s Honestly Significant Difference (HSD) test between each pair of stages is attached to the boxplot. All the comparisons are based on the log2-transformed RNA-Seq expressions (TPM-normalized). Before taking the log2 transformation, the zero expression of CCR3 or CCL24 is imputed by its none-zero minimum expression.

**Figure EV7. Drug treatment for sarcoidosis.**

(**A**) Images for the mice treated with vehicle and AZM daily for two weeks. (**B**) ELISA analysis of CCL24 for the sera of WT mice and KO mice administered by vehicle and AZM for over 24h. (**C**) Blood cell count for the WT mice and TSC KO mice that were administered by vehicle and CCL24/eotaxin-2 antibodies for 48h.

## References

Au - Ray A, Au - Dittel BN (2010) Isolation of Mouse Peritoneal Cavity Cells. JoVE: e1488

Au - Van Hoecke L, Au - Job ER, Au - Saelens X, Au - Roose K (2017) Bronchoalveolar Lavage of Murine Lungs to Analyze Inflammatory Cell Infiltration. JoVE: e55398

Bolger AM, Lohse M, Usadel B (2014) Trimmomatic: a flexible trimmer for Illumina sequence data. Bioinformatics 30: 2114–2120

Butler A, Hoffman P, Smibert P, Papalexi E, Satija R (2018) Integrating single-cell transcriptomic data across different conditions, technologies, and species. Nat Biotechnol 36: 411–420

Co DO, Hogan LH, Il-Kim S, Sandor M (2004) T cell contributions to the different phases of granuloma formation. Immunology Letters 92: 135–142

Comas M, Toshkov I Fau - Kuropatwinski KK, Kuropatwinski Kk Fau - Chernova OB, Chernova Ob Fau - Polinsky A, Polinsky A Fau - Blagosklonny MV, Blagosklonny Mv Fau - Gudkov AV, Gudkov Av Fau - Antoch MP, Antoch MP (2012) New nanoformulation of rapamycin Rapatar extends lifespan in homozygous p53-/- mice by delaying carcinogenesis. Aging

Costabel U, Hunninghake GW (1999) ATS/ERS/WASOG statement on sarcoidosis. Sarcoidosis Statement Committee. American Thoracic Society. European Respiratory Society. World Association for Sarcoidosis and Other Granulomatous Disorders. European Respiratory Journal 14: 735

Drent M, Crouser ED, Grunewald J (2021) Challenges of Sarcoidosis and Its Management. New England Journal of Medicine 385: 1018–1032

Elliott DC, Landon WL, Mark WJ, Sabahattin B, Wolfgang S, Peter W, Larry SS (2021) Phagosome-regulated mTOR signalling during sarcoidosis granuloma biogenesis. European Respiratory Journal 57: 2002695

Finak G MA, Yajima M, Deng J, Gersuk V, Shalek AK, Slichter CK, Miller HW, McElrath MJ, Prlic M, Linsley PS and Gottardo R (2015) MAST: a flexible statistical framework for assessing transcriptional changes and characterizing heterogeneity in single-cell RNA sequencing data. Genome Biol: 278

Fong K-w, Zhao JC, Lu X, Kim J, Piunti A, Shilatifard A, Yu J (2022) PALI1 promotes tumor growth through competitive recruitment of PRC2 to G9A-target chromatin for dual epigenetic silencing. Molecular Cell 82: 4611–4626.e4617

Fraser SD, Thackray-Nocera S, Shepherd M, Flockton R, Wright C, Sheedy W, Brindle K, Morice AH, Kaye PM, Crooks MG et al (2020) Azithromycin for sarcoidosis cough: an open-label exploratory clinical trial. ERJ Open Res 6

Fraser SD, Thackray-Nocera S, Wright C, Flockton R, James SR, Crooks MG, Kaye PM, Hart SP (2024) Effects of Azithromycin on Blood Inflammatory Gene Expression and Cytokine Production in Sarcoidosis. Lung 202: 683–693

Giovinale M, Fonnesu C, Soriano A, Cerquaglia C, Curigliano V, Verrecchia E, De Socio G, Gasbarrini G, Manna R (2009) Atypical sarcoidosis: case reports and review of the literature. Eur Rev Med Pharmacol Sci 13 Suppl 1: 37–44

Grunewald J, Grutters JC, Arkema EV, Saketkoo LA, Moller DR, Muller-Quernheim J (2019) Sarcoidosis. Nat Rev Dis Primers 5: 45

Grutters JC, Bosch JMMvd (2006) Corticosteroid treatment in sarcoidosis. European Respiratory Journal 28: 627–636

Haydar D, Cory TJ, Birket SE, Murphy BS, Pennypacker KR, Sinai AP, Feola DJ (2019) Azithromycin Polarizes Macrophages to an M2 Phenotype via Inhibition of the STAT1 and NF-κB Signaling Pathways. The Journal of Immunology 203: 1021–1030

Henske EP, Jóźwiak S, Kingswood JC, Sampson JR, Thiele EA (2016) Tuberous sclerosis complex. Nature Reviews Disease Primers 2: 16035

Hiranuma R, Sato R, Yamaguchi K, Nakamizo S, Asano K, Shibata T, Fukui R, Furukawa Y, Kabashima K, Miyake K (2024) Aberrant monocytopoiesis drives granuloma development in sarcoidosis. Int Immunol 36: 183–196

House NS, Welsh Jp Fau - English JC, 3rd, English JC, 3rd (2012) Sarcoidosis-induced alopecia. Dermatol Online J

Inoki K, Li Y Fau - Xu T, Xu T Fau - Guan K-L, Guan KL (2003) Rheb GTPase is a direct target of TSC2 GAP activity and regulates mTOR signaling. Genes Dev 17: 1829–1834

Jin S, Guerrero-Juarez CF, Zhang L, Chang I, Ramos R, Kuan C-H, Myung P, Plikus MV, Nie Q (2021) Inference and analysis of cell-cell communication using CellChat. Nature Communications 12: 1088

Joanne CM, Eóin NM, Lindsay H, Kelley EC, Joseph R, Sophie AF, Alfred DD, Holger KE, Anil KR, Cheryl AP et al (2014) Local hypersensitivity reaction in transgenic mice with squamous epithelial IL-5 overexpression provides a novel model of eosinophilic oesophagitis. Gut 63: 43

Judson MA, Chaudhry H, Louis A, Lee K, Yucel R (2015) The effect of corticosteroids on quality of life in a sarcoidosis clinic: The results of a propensity analysis. Respiratory Medicine 109: 526–531

Kawano S, Kato J, Kawano N, Yoshimura Y, Masuyama H, Fukunaga T, Shimao Y, Mihara K, Ueda A, Toyoda K et al (2012) Sarcoidosis manifesting as cardiac sarcoidosis and massive splenomegaly. Intern Med 51: 65–69

Kikuchi M, Koizumi A, Namisaki T, Asada S, Oyama M, Tomooka F, Fujimoto Y, Kitagawa K, Kawaratani H, Yoshiji H (2024) Improvement of liver histology in hepatic sarcoidosis due to treatment with corticosteroids and ursodeoxycholic acid: a case report. Clin J Gastroenterol 17: 327–333

Korsunsky I, Millard N, Fan J, Slowikowski K, Zhang F, Wei K, Baglaenko Y, Brenner M, Loh P-r, Raychaudhuri S (2019) Fast, sensitive and accurate integration of single-cell data with Harmony. Nature Methods 16: 1289–1296

Kournoutou GG, Dinos G (2022) Azithromycin through the Lens of the COVID-19 Treatment. Antibiotics doi: 10.3390/antibiotics11081063 [PREPRINT]

Krausgruber T, Redl A, Barreca D, Doberer K, Romanovskaia D, Dobnikar L, Guarini M, Unterluggauer L, Kleissl L, Atzmuller D et al (2023a) Single-cell and spatial transcriptomics reveal aberrant lymphoid developmental programs driving granuloma formation. Immunity 56: 289–306 e287

Krausgruber T, Redl A, Barreca D, Doberer K, Romanovskaia D, Dobnikar L, Guarini M, Unterluggauer L, Kleissl L, Atzmüller D et al (2023b) Single-cell and spatial transcriptomics reveal aberrant lymphoid developmental programs driving granuloma formation. Immunity 56: 289–306.e287

Kyriakos Saad M, El Hajj I, Saikaly E (2024) Sarcoidosis presenting as isolated massive splenomegaly: A case report. Clin Case Rep 12: e9283

Li B, Dewey CN (2011) RSEM: accurate transcript quantification from RNA-Seq data with or without a reference genome. BMC Bioinformatics 12: 323

Linke M (2017) Chronic signaling via the metabolic checkpoint kinase mTORC1 induces macrophage granuloma formation and marks sarcoidosis progression. Nat Immunol

Linke M, Pham HT, Katholnig K, Schnoller T, Miller A, Demel F, Schutz B, Rosner M, Kovacic B, Sukhbaatar N et al (2017) Chronic signaling via the metabolic checkpoint kinase mTORC1 induces macrophage granuloma formation and marks sarcoidosis progression. Nat Immunol 18: 293–302

Love MI, Huber W, Anders S (2014) Moderated estimation of fold change and dispersion for RNA-seq data with DESeq2. Genome Biology 15: 550

Lower EE, Smith Jt Fau - Martelo OJ, Martelo Oj Fau - Baughman RP, Baughman RP (1988) The anemia of sarcoidosis. Sarcoidosis 5: 51–55

Manzia (2011) Successful treatment of systemic de novo sarcoidosis with cyclosporine discontinuation and provision of rapamune after liver transplantation. Transpl Int

McGinnis CS, Murrow LM, Gartner ZJ (2019) DoubletFinder: Doublet Detection in Single-Cell RNA Sequencing Data Using Artificial Nearest Neighbors. Cell Systems 8: 329–337.e324

Megari K (2013) Quality of life in chronic disease patients. Health Psychology Research 1: e27

Meguro A, Ishihara M, Petrek M, Yamamoto K, Takeuchi M, Mrazek F, Kolek V, Benicka A, Yamane T, Shibuya E et al (2020) Genetic control of CCL24, POR, and IL23R contributes to the pathogenesis of sarcoidosis. Commun Biol 3: 465

Menzies-Gow A, Ying S, Sabroe I, Stubbs VL, Soler D, Williams TJ, Kay AB (2002) Eotaxin (CCL11) and Eotaxin-2 (CCL24) Induce Recruitment of Eosinophils, Basophils, Neutrophils, and Macrophages As Well As Features of Early- and Late-Phase Allergic Reactions Following Cutaneous Injection in Human Atopic and Nonatopic Volunteers1. The Journal of Immunology 169: 2712–2718

Mukhopadhyay S, Farver CF, Vaszar LT, Dempsey OJ, Popper HH, Mani H, Capelozzi VL, Fukuoka J, Kerr KM, Zeren EH et al (2012) Causes of pulmonary granulomas: a retrospective study of 500 cases from seven countries. Journal of Clinical Pathology 65: 51–57

Ng T, Yeghen T, Pagliuca A, Gillett DS, Mufti GJ (2000) Non-Caseating Granulomata Associated with Hypocellular Myelodysplastic Syndrome. Leukemia & Lymphoma 39: 397–403

Paolo C, Laura B, Francesco B, Maria Antonietta M, Piersante S, Paola R, Elena B (2019) Biomarkers of Sarcoidosis: a comparative study of serum chitotriosidase, ACE, lysozyme and KL-6. European Respiratory Journal 54: PA1959

Paus R, Maurer M, Slominski A, Czarnetzki BM (1994) Mast Cell Involvement in Murine Hair Growth. Developmental Biology 163: 230–240

Robinson MD, McCarthy DJ, Smyth GK (2009) edgeR: a Bioconductor package for differential expression analysis of digital gene expression data. Bioinformatics 26: 139–140

Sahin O, Ziaei A, Karaismailoğlu E, Taheri N (2016) The serum angiotensin converting enzyme and lysozyme levels in patients with ocular involvement of autoimmune and infectious diseases. BMC Ophthalmology 16: 19

Saito S, Kodama K, Kogiso T, Yamanashi Y, Taniai M, Ariizumi S, Yamamoto M, Tokushige K (2020) Atypical Sarcoidosis Diagnosed by Massive Splenomegaly. Intern Med 59: 641–648

Samiksha G, Miloni P, Rana Prathap P, Agam B, Salim D (2022) Role of Serum Soluble Interleukin-2 Receptor Level in the Diagnosis of Sarcoidosis: A Systematic Review and Meta-Analysis. medRxiv: 2022.2007.2016.22277713

Saxton RA, Sabatini DM (2017) mTOR Signaling in Growth, Metabolism, and Disease. Cell 168: 960–976

Shen Y, Pang C, Wu Y, Li D, Wan C, Liao Z, Yang T, Chen L, Wen F (2016) Diagnostic Performance of Bronchoalveolar Lavage Fluid CD4/CD8 Ratio for Sarcoidosis: A Meta-analysis. EBioMedicine 8: 302–308

Stuart T BA, Hoffman P, Hafemeister C, Papalexi E, Mauck WM, 3rd, Hao Y, Stoeckius M, Smibert P and Satija R (2019) Comprehensive Integration of Single-Cell Data. Cell: 1888–1902.e1821

Stuart T, Butler A, Hoffman P, Hafemeister C, Papalexi E, Mauck WM, Hao YH, Stoeckius M, Smibert P, Satija R (2019) Comprehensive Integration of Single-Cell Data. Cell 177: 1888-+

Subramanian A (2005) Gene set enrichment analysis: A knowledge-based approach for interpreting genome-wide expression profiles. PNAS

Tomita H, Sato S, Matsuda R, Sugiura Y, Kawaguchi H, Niimi T, Yoshida S, Morishita M (1999) Serum Lysozyme Levels and Clinical Features of Sarcoidosis. Lung 177: 161–167

Vukmirovic M, Yan X, Gibson KF, Gulati M, Schupp JC, DeIuliis G, Adams TS, Hu B, Mihaljinec A, Woolard TN et al (2021) Transcriptomics of bronchoalveolar lavage cells identifies new molecular endotypes of sarcoidosis. Eur Respir J 58

Weichhart T, Costantino G, Poglitsch M, Rosner M, Zeyda M, Stuhlmeier KM, Kolbe T, Stulnig TM, Horl WH, Hengstschlager M et al (2008) The TSC-mTOR signaling pathway regulates the innate inflammatory response. Immunity 29: 565–577

Xu J (2005) Preparation, Culture, and Immortalization of Mouse Embryonic Fibroblasts. Current Protocols in Molecular Biology 70: 28.21.21–28.21.28

Zhang H, Costabel U, Dai H (2021) The Role of Diverse Immune Cells in Sarcoidosis. Front Immunol

Zhu L (2014) TSC1 controls macrophage polarization to prevent injlammatory disease. Nat Commun

